# A Multiscale Framework for Uncovering Surfactant Mediated Viral Capsid Disruption

**DOI:** 10.1101/2025.06.16.659739

**Authors:** Srdan Masirevic, Jan K. Marzinek, Marcella L. Y. Kong, Grace Lin, Hongling Chen, Jiquan Liu, ChunSong Chua, C. Mark Maupin, Chandra S. Verma, Stephen J. Fox, Peter J. Bond

## Abstract

Disinfection remains a critical strategy for controlling the transmission of infectious diseases. However, small non-enveloped viruses exhibit exceptional resistance to many disinfectants, often requiring harsh protein-disrupting chemicals for effective inactivation, thereby limiting their applicability in personal care products due to associated side effects. Sodium dodecyl sulphate (SDS) is a widely used anionic surfactant known for its virucidal efficacy; however, the molecular details of its action against robust non-enveloped viruses remain poorly understood, limiting efforts to design safer and more targeted antiviral formulations. In this study, a multiscale simulation approach combining a novel atomic-resolution icosahedral “scaffold framework” and coarse-grained modelling was developed to elucidate the mechanism of SDS-driven disruption of MS2 bacteriophage capsid, a surrogate for non-enveloped viruses. Experimental analyses including dynamic light scattering and transmission electron microscopy revealed that SDS inactivates MS2 in a strongly pH-dependent manner, triggering capsid disassembly at acidic pH while leaving particles largely intact at neutral pH. Molecular dynamics simulations demonstrated that SDS micelles preferentially associate with hexameric pores and inter-dimer clefts under acidic conditions, where protonation of acidic residues weakens the electrostatic network of the capsid surface. Together, these findings provide a detailed molecular framework for SDS virucidal action and highlight the importance of environmental pH in modulating surfactant–virus interactions. These insights offer a foundation for designing next-generation antiviral surfactants with improved efficacy and biocompatibility.

## 1. Introduction

The ongoing threat of viral transmission in healthcare, food safety, and environmental contexts continues to underscore the critical need for effective antiviral interventions^1^. Sanitizing agents, such as alcohols, anionic surfactants, and quaternary ammonium compounds have been successfully used in commercially available formulations that have demonstrated virucidal properties to inactivate enveloped viruses in suspension and on surfaces^2^. Unlike enveloped viruses, which contain a lipid envelope and are generally more susceptible to membrane-disrupting agents, non-enveloped viruses rely on robust protein capsids for environmental persistence, making them considerably more difficult to inactivate^3^. In particular, human norovirus, a leading cause of viral gastroenteritis globally, poses a significant public health concern due to its extreme resistance to traditional chemical disinfectants, high heat stability, and low infectious dose^4–7^. As a result, sanitizing agents often exhibit limited efficacy against these pathogens, particularly under suboptimal application conditions^8–10^.

Among chemical disinfectants, surfactants have emerged as promising antiviral agents due to their unique amphiphilic properties, enabling them to disrupt biological membranes and protein assemblies effectively^11,12^. SDS, a widely used anionic surfactant, has demonstrated consistent antiviral activity against both enveloped and non-enveloped viruses by inducing protein denaturation and structural destabilization^13,14^. Despite its broad virucidal capabilities, the molecular basis underlying SDS-induced viral inactivation, particularly for non-enveloped viruses, remains incompletely understood. While early hypotheses attributed SDS activity to generalized protein denaturation or non-specific viral membrane disruption^15^, these explanations fall short in accounting for SDS’s highly variable efficacy under different environmental conditions—most notably, the strong influence of pH^16,17^. Previous studies have shown that SDS exhibits greater virucidal potency against non-enveloped viruses at acidic pH, suggesting that its mechanism likely involves specific structural interactions with viral protein capsid components that are modulated by charge distribution and protonation states^14^. Moreover, protein denaturation alone does not fully explain the selective disruption of robust icosahedral capsids, raising the possibility that SDS may preferentially bind to and destabilize key interfacial regions within the viral architecture. Therefore, a detailed mechanistic understanding of how SDS interacts with specific capsid elements, and how environmental factors like pH modulate this process, are essential for rational design of next-generation antiviral surfactants.

The MS2 bacteriophage, a small single-stranded RNA (+ssRNA) virus with a well-characterized icosahedral capsid, is a frequently used surrogate for human enteric viruses in disinfection studies because of its environmental relevance, high infectivity, and structural similarity to pathogenic species, such as norovirus^18^. The MS2 capsid, measuring ∼23–28 nm in diameter, comprises one maturation protein (MP) and 178 capsid protein (CP) units. Each CP unit consists of 129 amino acids arranged into an interior-facing five-stranded β-sheet and an exterior hairpin structure with two helices. The capsid is built from CP dimers in two forms: symmetric (C/C) dimers at the 2-fold axes, and asymmetric (A/B) dimers at the 3-fold and 5-fold axes. This structural variation allows the MS2 capsid to adopt its characteristic icosahedral geometry. While the ability of SDS to inactivate MS2 is known^19^, the exact structural vulnerabilities that render the virus susceptible in such environments remain poorly defined. This raises a key mechanistic question: how does environmental pH influence SDS access, binding, and structural impact within the viral architecture?

Previous all-atom MD simulations have provided high-resolution insights into MS2 stability and dynamics, highlighting key structural components such as dimer interfaces and pore regions that contribute to its remarkable robustness^20–22^. However, the intrinsic limitations of all-atom approaches—particularly their computational expense and inability to access biologically relevant timescales, particularly for large systems such as entire viral capsids— pose major challenges in studying large-scale destabilization events induced by surfactants such as SDS. Capturing events that may be mechanistically important such as micelle formation, surface binding, capsid pore expansion, or global capsid rearrangements requires both longer simulation times and larger system sizes that are typically not feasible to study at atomic resolution. To overcome these limitations, in this study we report a coarse-grained (CG) MD simulation of the MS2 capsid, structurally calibrated against an all-atom model, and then apply this to investigate the full capsid-scale dynamics of SDS-mediated disruptions. In a complementary approach, a simplified “scaffold model” was developed for all-atom simulations, focusing specifically on hexameric and pentameric pores surrounded by adjacent dimers. This reduced model provided a computationally feasible yet experimentally verifiable platform for directly validating key observations from the coarse-grained simulations without modelling the entire capsid. Supporting experimental methodologies, including dynamic light scattering (DLS) and transmission electron microscopy (TEM), were utilized to confirm structural changes and disruption of the MS2 capsid under acidic conditions, thereby strengthening the validity of our computational findings.

Building on this integrated multiscale approach, elucidation of the molecular basis for SDS-mediated capsid disruption, focusing on the dynamics, critical binding sites, and structural transitions that govern susceptibility to surfactant-induced destabilization was evaluated. By doing so, a deeper mechanistic understanding is provided on how surfactant properties and environmental factors converge to promote disassembly of non-enveloped viruses, which should ultimately help to guide the rational design of safer, more effective antiviral strategies.

## 2. Materials and Methods

### 2.1 Experimental Procedures

#### 2.1.1 Bacteria host and bacteriophage propagation

*Escherichia coli* strain C3000 (ATCC® 15597) was used as the host for *Escherichia coli* bacteriophage MS2 (ATCC® 15597B1). The bacterial strain was initially streaked on Luria-Bertani (LB) agar and incubated at 37□°C for 18–24 hours. A single colony was then subcultured in 20□mL of LB broth and incubated at 37□°C with agitation (shaking) for another 18–24 hours. Subsequently, 100□µL of this overnight culture was inoculated into fresh 20□mL LB broth and incubated at 37□°C with agitation for 4 hours to reach the logarithmic growth phase. This culture was used as the working bacterial stock for downstream applications.

MS2 phage was propagated using the log-phase *E. coli* C3000 host. A single plaque of MS2 was picked, resuspended in 1□mL of LB broth, and centrifuged at 12,000□rpm for 5 minutes to clarify. The supernatant containing the phage was mixed with 1□mL of log-phase *E. coli* C3000 and 4□mL of 0.75% (w/v) LB soft agar, then overlaid on LB agar plates. The plates were incubated overnight at 37□°C. The next day, LB broth was added to the plate and gently swirled for 15 minutes at room temperature to release the phage particles. The resulting suspension was collected and centrifuged at 12,000□rpm for 20 minutes to remove bacterial debris. The clarified phage-containing supernatant was stored at 4□°C until further use.

#### 2.1.2 Virucidal efficacy testing against MS2

A total of 800□µL of SDS solution was mixed with 100□µL of 3% bovine serum albumin (BSA) to achieve a final BSA concentration of 0.3%, followed by the addition of 100□µL of MS2 bacteriophage suspension (1□×□10□□PFU/mL). The reaction mixture was incubated at room temperature for the designated contact time. After incubation, 10-fold serial dilutions were prepared in LB broth. Subsequently, 50□µL of each dilution was plated in triplicates onto *E. coli* strain C3000 overlays, which consisted of log-phase *E. coli* C3000 diluted 10-fold and embedded in 0.75% (w/v) LB soft agar, overlaid onto LB agar plates. The plates were incubated at 37□°C for 18–24 hours, after which plaque-forming units (PFU) were enumerated and expressed as PFU/mL.

#### 2.1.3 DLS analysis

Concentrated and purified MS2 bacteriophage was first diluted 10-fold in TNE buffer (pH 8.0), followed by filtration through a 0.2□µm membrane to remove viral aggregates. Subsequently, 20□µL of the MS2 suspension was added to 80□µL of the test sample and incubated at room temperature for 5 minutes. After incubation, 80□µL of the reaction mixture was transferred into a disposable cuvette for analysis. Particle size measurements were performed at 25□°C using a Malvern Zetasizer Nano-ZS instrument (ZEN3600, Malvern Instruments, Worcestershire, UK), equipped with a 4□mW He-Ne laser (λ = 633□nm), at a backscattering detection angle of 173°. A clear disposable capillary cell (DTS0012) was used for the measurements. The size distribution was obtained by deconvolution of the measured intensity autocorrelation function using a non-negatively constrained least-squares fitting algorithm. Each sample was measured in triplicate.

#### 2.1.4 TEM analysis

Negative-staining TEM images were acquired using a JEM 1400Plus transmission electron microscope (JEOL) operating at an accelerating voltage of 120□kV. Samples were prepared by depositing a drop of the sample solution onto a carbon-coated copper grid and allowing it to adsorb for 5 minutes. The grid was then washed and negatively stained with 2% (w/v) uranyl acetate for 2 minutes. Grids were air-dried prior to imaging.

### 2.2 Computational Modelling - Methods and Analysis

#### 2.2.1 CG simulation – system setup

An all-atom model of the MS2 bacteriophage capsid was obtained from the Protein Data Bank (PDB ID: 2MS2)^23^. The all-atom structure was converted to a CG representation using the Martini 2.2 force field^24,25^. CG parameters for SDS molecules were adopted from the standard Martini 2.2 force field without additional modifications. The MP was excluded from the simulation systems to isolate and focus on capsid-specific interactions and structural perturbations by SDS; additional details and validation analyses comparing simulations with and without MP are provided in the *Supplementary Materials*. The simulation box was cubic with dimensions of 32 × 32 × 32□nm³. As the viral RNA was not explicitly included in the CG model, 3,600 chloride ions were placed within the capsid to approximate the electrostatically negative environment typically provided by the encapsulated genome. The solvent environment consisted of ∼173,600 standard Martini water beads (90%) and 10% antifreeze beads, with the addition of Na^+^ and Cl^−^ ions to reach an ionic concentration of 150 mM. Energy minimization was performed using the steepest descent (SD) algorithm for 5,000 steps with a maximum step size of 0.01□nm and a force convergence criterion of 1,000□kJ mol□¹□nm□¹. Subsequently, the system was equilibrated for the total of 100 ns, comprising initial NVT, followed by NPT ensemble simulation with position restraints on protein backbone beads.

To maintain native-like structural integrity of the MS2 capsid while allowing flexibility, an elastic network (EN) was applied to backbone beads within 0.5-0.9 nm cutoff distance in each dimer of structured regions, excluding loops. Multiple EN force constants (500–10,000 kJ mol^−1^ nm^−2^) were tested to determine optimal stability by comparing the backbone RMSD profiles of CG models with reference all-atom simulations (*Supplementary materials*). The final production simulations used an optimized force constant of 3,000 kJ mol^−1^ nm^−2^. An assessment of Martini 3 was also conducted; however, the MS2 capsid displayed pronounced structural instabilities, and was thus excluded from further study (*Supplementary materials*).

#### 2.2.2 CG simulation - simulation details

All CG MD simulations were performed using GROMACS 2018.3 simulation package^26^. Temperature and pressure were maintained at 310 K and 1 bar using a velocity-rescaling thermostat (τ = 1.0 ps) and a Parrinello–Rahman barostat^27^ (τ = 12.0 ps), respectively, following standard protocols recommended for Martini coarse-grained simulations. Bond lengths were constrained using the LINCS algorithm, and equations of motion were integrated using the leap-frog integrator with a 20 fs timestep. Nonbonded interactions were calculated using a cutoff scheme with a cutoff distance of 1.1 nm for both Lennard-Jones and Coulomb interactions. Electrostatics were computed using the reaction-field method with a dielectric constant of 15, consistent with the Martini 2.2 force field. Simulations were performed on: Procter & Gamble internal high-performance computing cluster, as well as on Singapore’s National Supercomputing Center (https://www.nscc.sg) using up to 24 nodes containing 24 CPUs each (Intel® Xeon® CPU E5-2960 v3 @ 2.60.

To examine detergent effects, 10 wt. % SDS by solvent weight was introduced by calculating the required number of SDS molecules based on total water bead mass. Simulations were run for 10 µs per condition (pH 2.5 and 7.5), both with and without SDS, under the NPT ensemble. The pH 2.5 and pH 7.5 models were generated by modifying the protonation states of titratable residues based on standard pKa values, with acidic residues: aspartic acidic (Asp), and glutamic acid (Glu) protonated at pH 2.5 and deprotonated at pH 7.5.

#### 2.2.3 CG simulation - Analysis

All trajectory analyses were performed using MDAnalysis and in-house Python scripts unless otherwise specified^28^. To examine electrostatic contributions to capsid stability, salt bridges were calculated between Asp and Glu, and basic lysine (Lys), and arginine (Arg) residues using a 5□Å distance cutoff, averaged over the final 2 μs of each trajectory. To localize SDS–capsid interactions, SDS density and occupancy maps were generated using GROMACS analysis tools to identify high-frequency SDS binding regions. Residue-level SDS contact frequencies were calculated across the trajectory and normalized by the total number of residues of each type in the MS2 capsid to correct for composition bias. Center of mass (COM) distances between selected dimer pairs were computed to monitor SDS-induced dimer separation. Tilt angles were calculated for individual dimers based on the change in orientation of their principal axes relative to the starting frame. The radius of gyration (Rg) was measured for each dimer in selected hexamers to quantify internal structural compactness and flexibility under different simulation conditions. To assess SDS penetration depth, the radial distances between each SDS headgroup and the capsid COM were computed and compared to the radial span of the capsid itself (i.e., distance from the capsid COM to the COMs of the innermost and outermost capsid protein beads). This enabled classification of SDS molecules as “internal” or “external” based on their relative radial positions. Pore radius measurements were performed using a center-based approach. First, loop residues (residue numbers) from each monomer forming the pore were selected to define the pore perimeter. The COM of these loop residues was calculated for each monomer in every frame. Next, the overall pore center was determined as the COM of all selected monomer loop COMs. Finally, the pore radius was computed as the average distance between the pore center and each monomer loop COM across the trajectory. Pore diameters were then averaged over time and categorized based on SDS occupancy and pH condition. Where appropriate, statistical comparisons were performed using Welch’s t-test^29^, and error bars represent standard deviations.

### 2.3 All atom simulation system setup

#### 2.3.1 Protein scaffold system setup

The experimental structure of MS2 (PDB ID: 2MS2) was used to create a minimal protein scaffold system that is solely focused on the pore regions which correspond to capsid icosahedral pentameric and hexameric units. A single hexamer or pentamer was extracted together with all surrounding protein dimers. The pentamer construct is referred to as the 5-fold axis that is composed of 5 dimers surrounding the pore. For the hexamer this corresponds to flat faces of the icosahedron between vertices (5-fold) and 2-fold edges (2-fold axis) composed of 6 dimers surrounding the pore. In total there were 20 protein monomers for the pentamer: 5 dimers for the pentamer and the protein scaffold made of 5 dimers surrounding the central pentamer. In the case of the hexamer the construct was composed of 6 dimers for the hexamer and protein scaffold made of 6 dimers surrounding the central hexamer. In each system protein charges were assigned according to neutral pH with charged termini using the CHARMM36m^30^ force field. Both systems were subject to energy minimization in vacuum using the SD algorithm with a 0.01 nm step size and 500 kJ mol^-1^ nm^-1^ force tolerance. Each construct was placed in a cubic box of ∼20x20x10 nm^3^. Approximately 110,000 TIP3P water^31^ molecules were added to the box along with 150 mM NaCl salt whilst neutralizing the overall system charge. Energy minimization was performed using the SD algorithm with the same settings. Each system was equilibrated in the NPT ensemble for 5 ns with position restraints on protein backbone atoms with a 1,000 kJ mol^-1^ nm^-2^ force constant. Production runs corresponded to 200 ns in the NPT ensemble and involved position restraints on protein backbone scaffold surrounding the pentamer or hexamer. Several force constants of the scaffold position restraints were tested which corresponded to: 10, 50, 200, 500 and 1,000 kJ mol^-1^ nm^-2^ and resulted in 5 independent simulations for either pentamer or hexamer.

#### 2.3.2 SDS interactions with pentamer/hexamer

According to experimental data, two pH conditions were studied: pH 5 and pH 3. Since no histidine residues are present in the MS2 sequence, the protein was treated as being at neutral pH with charged termini for the pH 5 condition. For the pH 3 condition, all Asp and Glu residues were modeled in their protonated forms. In both pH conditions, SDS was modeled in its negatively charged form (pKa ≈ 3.3). A total of four systems were studied: i) Hexamer at pH 3 with 5 wt% SDS; ii) Hexamer at pH 5 with 5 wt% SDS; iii) Pentamer at pH 3 with 5 wt% SDS; and iv) Pentamer at pH 5 with 5 wt% SDS.

Each protein construct, along with its corresponding protein scaffold (see previous section), was placed in a cuboidal simulation box of dimensions 16x16x15 nm³. To match the experimental SDS concentration of ∼5 wt%, 40 SDS molecules were added to each box and positioned above the outer surface of either the pentamer or hexamer. Each system was solvated with approximately 100,000 TIP3P water molecules and 150 mM salt, while ensuring overall system charge neutrality. To prevent SDS periodic interactions with the RNA-facing side of the protein, a water layer 0.6–0.8 nm thick was placed and maintained in a position-restrained state throughout equilibration and production runs. For these restraints, a force constant of 1,000 kJ mol□¹ nm□² was applied to the oxygen atoms of the restrained water molecules. The protein scaffold backbone atoms were restrained during both equilibration and production runs using an optimal force constant of 10 kJ mol□¹ nm□². Each system was equilibrated in the NPT ensemble for 10 ns, with additional position restraints applied to the pentamer/hexamer backbone atoms (1,000 kJ mol□¹ nm□²) and SDS heavy atoms. Production simulations were performed for 1 μs in the NPT ensemble, maintaining position restraints on the scaffold backbone surrounding the pentamer or hexamer, as well as the water layer described above.

#### 2.3.3 Simulation details

All simulations were performed using the GROMACS 2018.3 simulation package^26^. A temperature of 310 K was maintained using the velocity rescaling thermostat with an additional stochastic term using a time constant of 1 ps. Pressure was maintained isotropically at 1 atm using the Berendsen barostat^32^ during equilibration and Parrinello-Rahman barostat^33^ during production run and a time constant of 5 ps was used. All bonds which involved hydrogens were constrained using the LINCS algorithm. Equations of motion were integrated using the leap-frog algorithm with a time step of 1 fs during initial two stages of equilibration and 2 fs for the last part of equilibration and the production run. Long-range electrostatic interactions were described using the particle mesh Ewald method.^34^ The short-range electrostatics real space cut-off was 1.2 nm and the short-range van der Waals cut-off was also 1.2 nm. Periodic boundaries conditions were applied in all directions. Simulations were performed on: Singapore’s National Supercomputing Center (https://www.nscc.sg) using up to 24 nodes containing 24 CPUs each (Intel® Xeon® CPU E5-2960 v3 @ 2.60.

## 3. Results

### 3.1 SDS virucidal efficacy against MS2 – experimental study

SDS-induced inactivation of MS2 bacteriophage was strongly modulated by both pH and SDS concentration. At all tested concentrations (0.5%, 1%, 5%), SDS achieved near-complete viral inactivation at pH 2.5, with log reductions exceeding 5 logs. However, a modest increase in pH to 3.0 led to a notable decline in SDS efficacy, particularly at lower SDS concentrations. At pH 3.5, SDS lost almost all virucidal activity across all concentrations tested (**Figure 1**). These results reveal a sharp efficacy threshold governed by pH, where even a 0.5 unit increase above pH 2.5 can drastically reduce antiviral performance. This underscores the critical need for precise pH control in SDS-containing formulations to preserve their antiviral potency.

**Figure 1.**
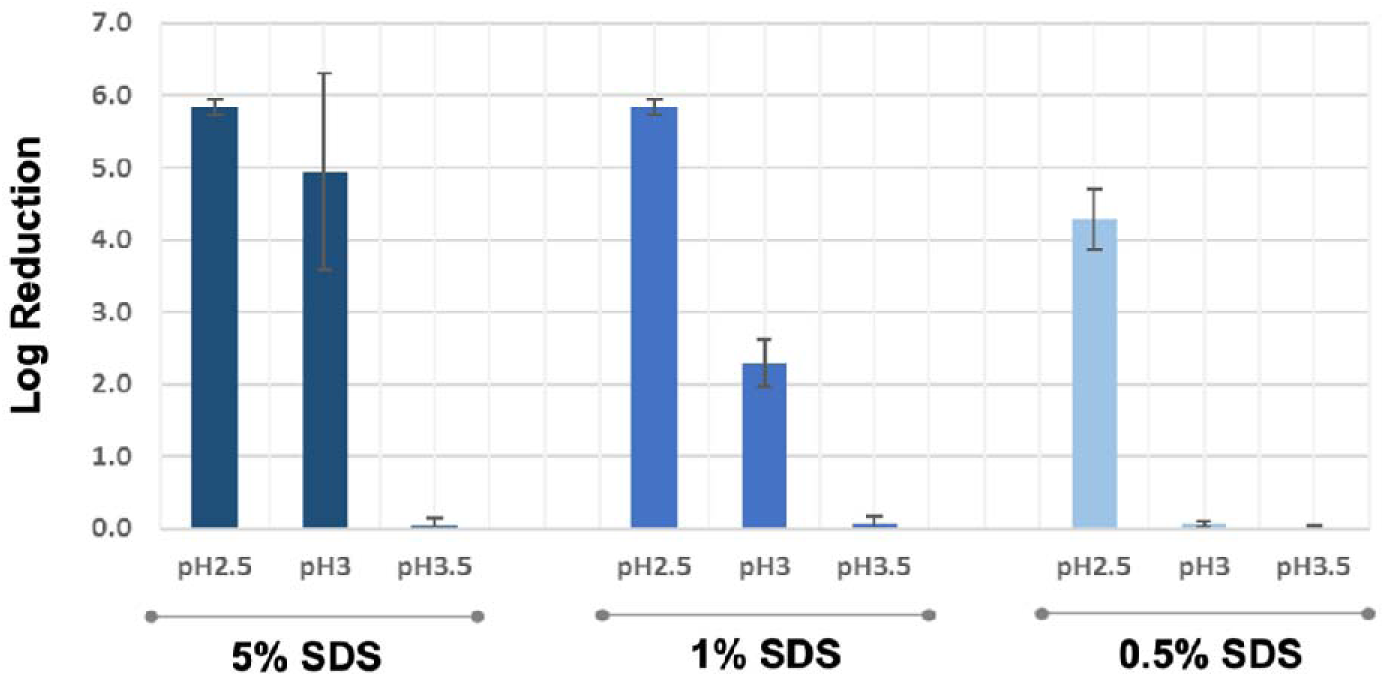
Virucidal efficacy of SDS at various concentrations and pH values against MS2 bacteriophage after 10 minutes of treatment. MS2 bacteriophage was exposed to SDS at 5%, 1%, and 0.5% concentrations across a pH range from approximately 2.5 to 3.5. The virucidal activity of SDS was found to be pH-dependent, with significantly higher efficacy observed at pH 2.5 across all concentrations tested. Data shown represent the mean ± standard deviation from two independent experiments.

TEM and DLS analyses provided complementary evidence regarding the effect of 1% SDS treatment on MS2 bacteriophage at pH 2.5 and pH 7.5. TEM imaging of untreated MS2 particles (pH 7.5) revealed well-defined individual virus particles as well as clustered aggregates (**Figure 2A**), consistent with the DLS observation of two distinct size populations. Following exposure to 1% SDS at pH 2.5, TEM images clearly demonstrated a substantial loss of intact viral particles (**Figure 2B**), indicating significant structural disruption or complete disassembly of the viral capsid under acidic SDS conditions. Correspondingly, DLS analysis revealed a marked decrease in the larger-size particle population, with a concurrent emergence of smaller-sized entities, likely attributable to SDS-induced micellar formation. Collectively, these findings highlight the destabilizing effect of SDS under acidic conditions and suggest a structural basis for its pH-dependent virucidal activity.

**Figure 2.**
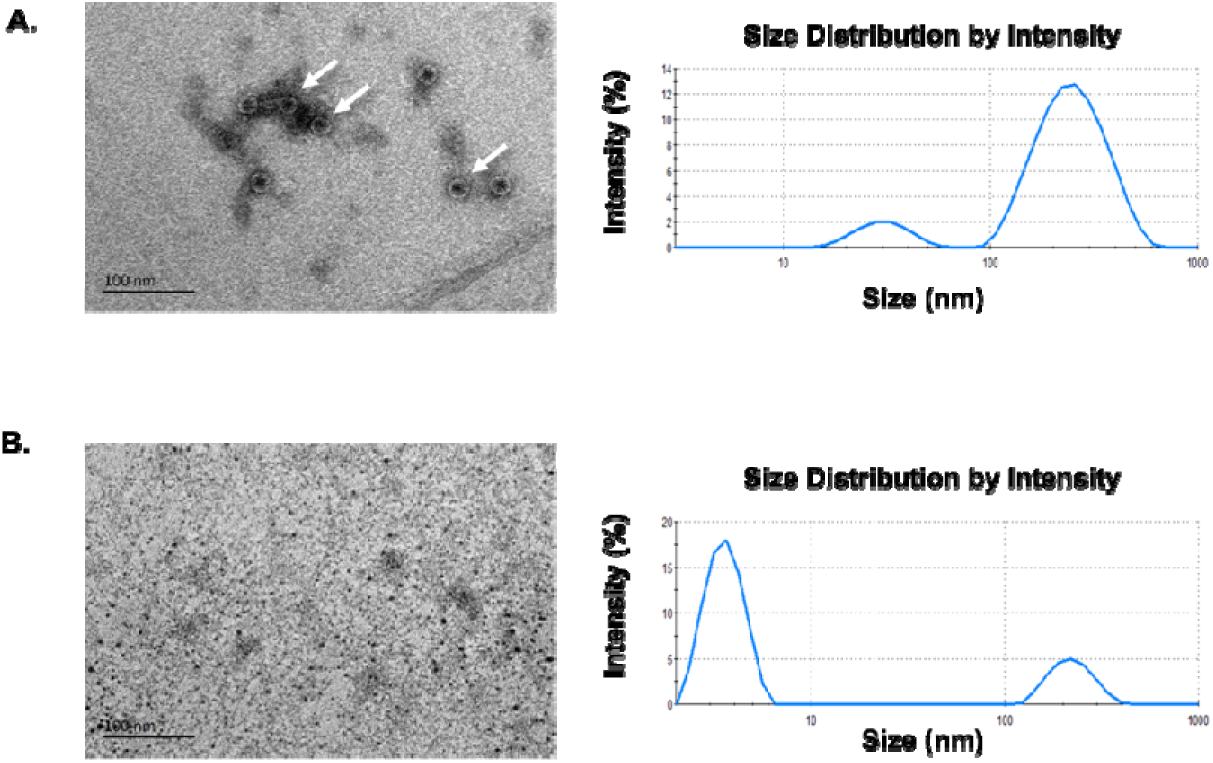
TEM and DLS analyses of MS2 following 1% SDS treatment at pH 2.5. **A.** TEM image of untreated MS2 at pH 7.5, showing both individual particles and clusters (indicated by arrows). DLS analysis revealed two size populations: a smaller peak likely corresponding to single MS2 particles and a larger peak representing clustered particles. **B.** TEM image of MS2 treated with 1% SDS at pH 2.5, showing a marked loss of intact bacteriophage particles. Corresponding DLS analysis demonstrated a reduction in the larger cluster-associated peak and the emergence of a smaller size peak (arrow), likely corresponding to SDS-generated micellar structures.

### 3.2 Coarse-grained simulations reveal pH-dependent and SDS-driven destabilization of the capsid

#### 3.2.1 Effect of pH on MS2 capsid stability

The current and previous^19^ experimental studies demonstrated distinct changes in MS2 capsid stability under varying pH conditions, potentially suggesting significant structural alterations and differential susceptibility to surfactants like SDS. Motivated by these observations, CG MD simulations in the absence of SDS were employed to further elucidate the mechanistic effects of pH variations at the molecular level. Visual inspection of the capsid structures at the end of the 10 µs simulations revealed more pronounced distortions at pH 2.5 (**Figure S2A**) compared to the well-preserved icosahedral shape observed at pH 7.5 (**Figure S2B**). To quantify these differences, salt bridge contacts formed between acidic (Glu, Asp) and basic (Lys, Arg) residues were analyzed over the final 2 μs of simulations. As shown in **Figure S2C**, the number of salt bridges was markedly lower at pH 2.5 (mean ≈ 1052) than at pH 7.5 (mean ≈ 1457), consistent with reduced electrostatic stabilization due to protonation of acidic residues under acidic conditions.

#### 3.2.2 SDS binding and destabilization of MS2 capsid

To uncover the molecular mechanism underlying SDS-induced MS2 capsid destabilization, CG simulations were performed at both acidic (pH 2.5) and neutral (pH 7.5) conditions using 10 wt. % SDS. SDS molecules rapidly self-assembled into spherical micelle-like aggregates with diameters of ∼3–5 nm (**Figure 3A**), consistent with the known aggregation behaviour of anionic surfactants at high concentrations. To quantify the spatial preferences of SDS micelles, occupancy analyses and lifetime measurements for SDS–capsid interactions were conducted over the 10□μs simulation time. As illustrated in **Figure 3B and 3C** SDS micelles preferentially associated with the dimer-dimer interfaces and hexameric and pentameric pores, where SDS clusters can be seen accumulating at inter-dimer clefts and pore openings. A similar number of total SDS–capsid binding events were observed at both pH 2.5 and pH 7.5, with broadly similar average interaction lifetimes (∼24–25 ns; **Figure S3A**). However, estimation of interactions between the negatively charged SDS headgroup and positively charged residues (Lys, Arg), resulted in a pH-dependent difference. The lifetime distributions revealed more frequent and longer-lived interactions at pH 2.5, in comparison to the pH 7.5 condition (**Figure S3B)**. Complementary residue-level contact analysis reinforced this interpretation (**Figure S4A**). Aromatic (Trp, Phe) and hydrophobic residues (Leu, Val, Pro) exhibited the strongest normalized contact frequencies with SDS across both pH conditions. Notably, Glu, Asp exhibited significantly more SDS interactions at pH 2.5 compared to pH 7.5. This is consistent with their protonated states at low pH reducing charge repulsion and enhancing SDS accessibility. Structural mapping of these residues (**Figure S4B**) revealed that the involved aromatic residues clustered at inter-dimer interfaces, while the hydrophobic and acidic residues were more prevalent at pore regions—hotspots that spatially align with the SDS binding zones observed in **Figure 3B and Figure 3C**.

**Figure 3.**
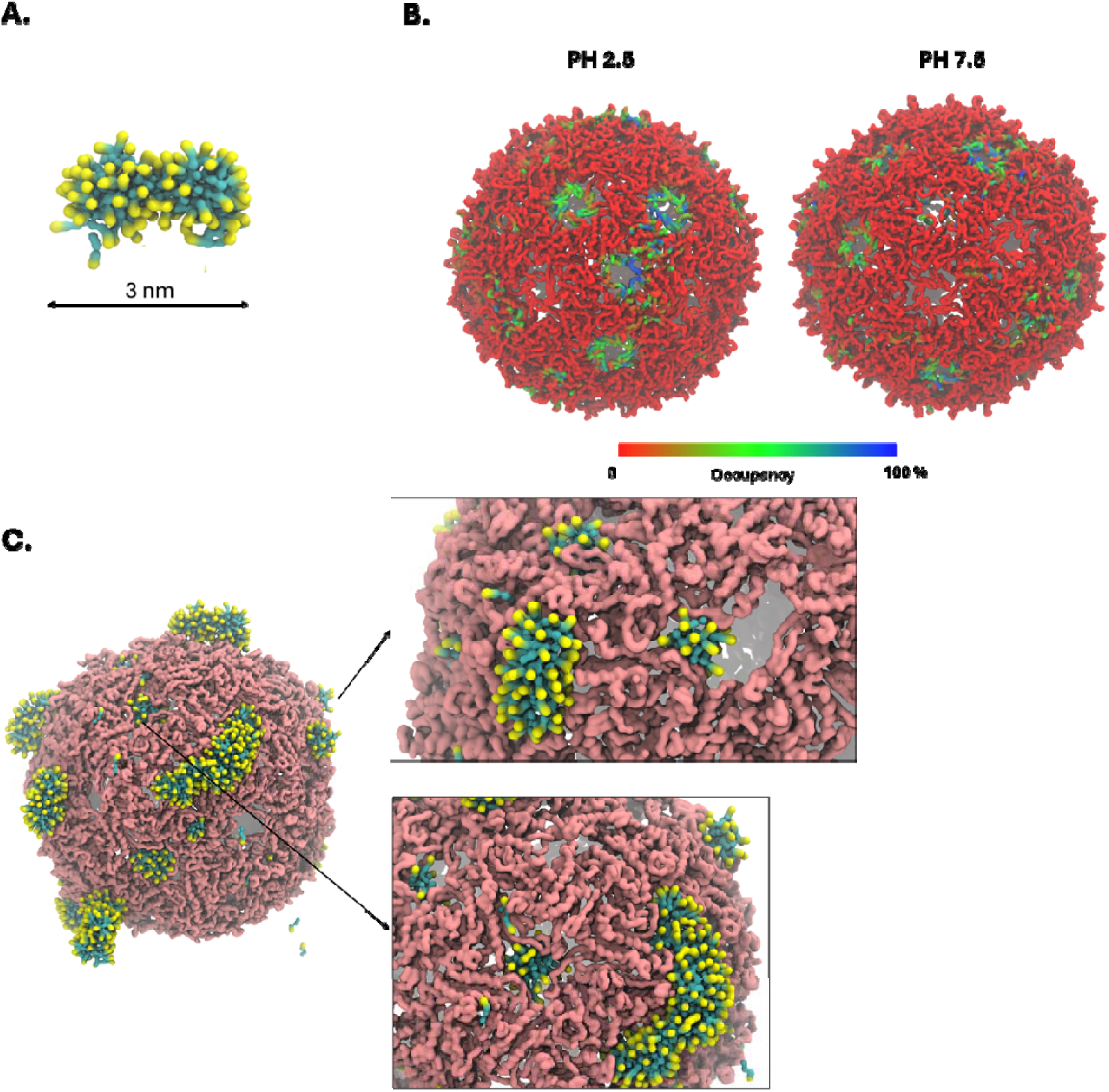
CG simulations reveal SDS micelle formation and capsid interaction hotspots. **A.** Representative micelle-like SDS aggregate formed at 10 wt. % SDS concentration, with a diameter of ∼3□nm. **B.** Surface occupancy maps showing SDS accumulation on the MS2 capsid at pH 2.5 (left) and pH 7.5 (right). Red regions denote low occupancy, blue regions denote high occupancy. **C.** Full capsid representation at pH 2.5 illustrating widespread SDS binding across the outer surface. SDS micelles (yellow – headgroup and blue - tail) are shown in association with key structural sites on the capsid backbone (pink).

**Figure 4.**
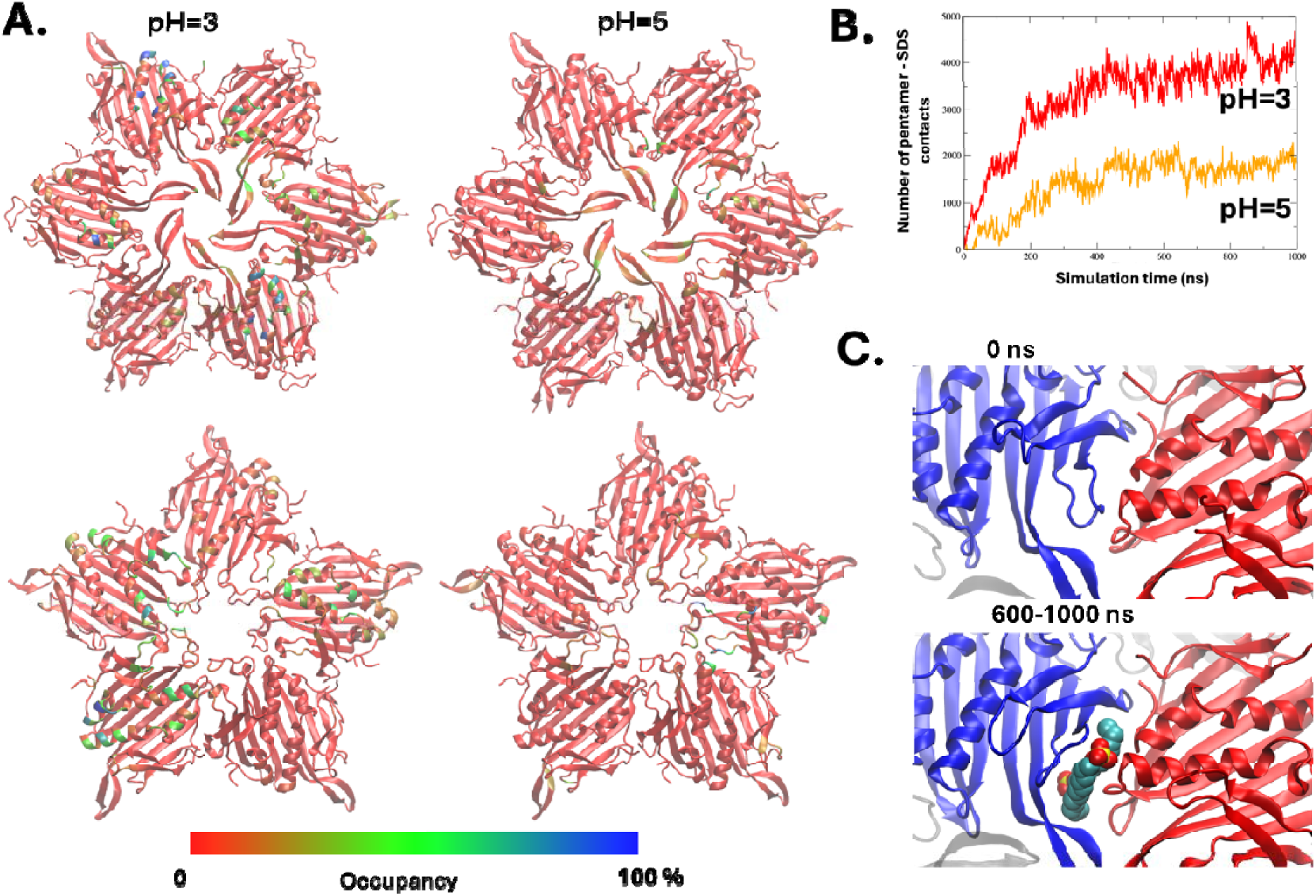
SDS binding hotspots revealed by all-atom simulations of MS2 hexameric scaffold. **A.** Snapshot of an all-atom simulation. **B.** Number of SDS contacts with the scaffold at pH 3 compared to pH 5 over the course of simulations. **C.** Zoomed-in view of a representative SDS-binding region at the interface of two adjacent dimers, showing detailed residue-level interactions.

Next, analysis of capsid pore diameters across pH 2.5 and pH 7.5 conditions revealed consistent differences between hexameric and pentameric pores in both SDS-treated and reference systems (SDS-free). These pore types correspond to geometrically distinct regions of the icosahedral capsid, with pentameric pores typically positioned at 5-fold vertices and hexameric pores located between them. At pH 2.5, hexameric pores exhibited higher SDS occupancy than pentameric pores. On average, 12 out of 20 hexameric pores (60%) were occupied, with 6 (30%) consistently hosting large SDS micelles (more than 10 SDS molecules). In contrast, pentameric pores showed 50% (6 out of 12) occupancy, with approximately 42% occupied by large micelles. At pH 7.5, the trend remained, with 10 out of 20 hexameric pores (50%) occupied on average, including 45% with large micelles, while pentameric pores averaged 42% occupancy, and only 25% accommodated large SDS clusters.

Comparing pore diameter changes, hexameric pores at pH 2.5 exhibited pronounced expansion when occupied by SDS micelles, increasing by an average of 3.7 Å from 37.4 Å in the reference systems to 41.0 Å in SDS-treated systems. Unoccupied hexameric pores showed minimal change, with diameters shifting from 37.3 Å to 38.0 Å. At pH 7.5, however, SDS binding did not lead to significant expansion. Occupied hexameric pores slightly contracted by 0.4 Å, while unoccupied pores expanded modestly by 0.8 Å. These trends suggest that SDS-induced widening of hexameric pores is strongly pH-dependent and more pronounced under acidic conditions, likely due to enhanced SDS aggregation and favourable surface interactions. For pentameric pores, the average expansion for SDS-occupied pores at pH 2.5 was 2.8 Å, with diameters increasing from 38.0 Å to 40.7 Å, while unoccupied pores expanded slightly by 1.2 Å. At pH 7.5, occupied pentameric pores increased in diameter by 2.1 Å, whereas unoccupied ones exhibited a slight contraction of 0.6 Å. Together, these results indicate that SDS micelle binding leads to pore expansion more consistently under acidic conditions, with hexameric pores showing a more substantial and pH-sensitive response than pentameric pores.

#### 3.2.3 SDS-induced structural perturbation of dimer interfaces at acidic pH

To investigate broader structural events upon SDS binding to hexameric pores, CG simulations were extended beyond the initial 10 µs in which SDS-bound hexameric pores at pH 2.5 showed a mean diameter increase of ∼3.7 Å as described above. To explore the longer-term effects of this expansion, a restrained scaffold model consisting of hexameric pore units surrounded by adjacent capsid protein dimers was constructed, and two coarse-grained 5 µs simulations at pH 2.5 were conducted: one where the SDS micelle was retained, and another where it was removed. Analysis of pore diameters over these extended trajectories revealed that the pore with the SDS micelle expanded further to an average of 45.3 Å, while the SDS-free counterpart remained at approximately 41.2 Å, consistent with its original expanded state (**Figure S5**).

Next, to further investigate structural perturbations at the dimer interface level, the COM distance distributions between selected dimer pairs within the selected hexamer (**Figure S6**) were evaluated. The most significant increase was observed between Dimer 5 and Dimer 6, where the SDS micelle was concentrated at the pore and dimer interface (**Figure S7**). Their average COM distance increased from 44.3 Å in the reference system to 46.1 Å in the SDS-bound system (Δ ≈ 1.8 Å). In contrast, other dimer pairs such as Dimer 4–5 and Dimer 1–6 exhibited smaller increases (∼0.5–1.7 Å), indicating smaller and localized structural displacement near the SDS micelle. To assess whether these structural shifts were accompanied by orientational changes, the tilt angles of Dimer 5 and Dimer 6 using their principal axes relative to their initial orientations were quantified. Under SDS-free conditions, the mean tilt angles were 3.7° (Dimer 5) and 2.3° (Dimer 6), with RMS deviations of 4.4° and 2.5°, respectively. In the SDS-bound system, mean tilt angles increased to 7.9° and 3.3°, and RMS deviations rose to 8.6° and 3.7° for Dimers 5 and 6, respectively. These results suggest that SDS promotes local rotational mobility or “wobbling” of the dimers under micelle exerted pressure. Next, the internal compactness of each dimer was examined using radius of gyration (Rg) analysis (**Figure S8)**. Dimer 5, which was in closest contact with the SDS micelle, displayed the most pronounced increase in Rg under SDS, indicating a possible structural loosening of its structure. Other dimers exhibited varying but generally consistent increases in Rg, further supporting the idea of SDS-induced destabilization and expansion. Finally, radial distribution analysis of SDS headgroups relative to the capsid COM revealed distinct pH-dependent localization patterns (**Figure S9**). At pH 2.5, 76.8% of the SDS molecules were located within the outer capsid boundary, compared to only 44.0% at pH 7.5, indicating significantly deeper penetration of SDS into the capsid structure under acidic conditions, which likely contributes to enhanced destabilization and disassembly at low pH.

### 3.3 All-atom simulations **–** SDS binding patterns and surface charge effects at different pH conditions

To determine whether the SDS binding preferences and destabilization patterns observed in CG simulations are also reproduced at atomic resolution, all-atom MD simulations were performed using our simplified scaffold model. Given the prohibitive computational cost of simulating the entire MS2 capsid at all-atom resolution over biologically relevant timescales^22^, a representative model consisting of hexameric and pentameric pore units flanked by adjacent capsid protein scaffold made of dimers at each side was constructed (see Methods). This approach allowed us to capture local binding interactions and surface-specific electrostatic changes with high precision, while maintaining key structural features critical for SDS–capsid association. Analysis of SDS occupancy revealed that at both pH conditions, SDS molecules preferentially accumulated at pore entrances and inter-dimer interfaces (**Figure 3A**). However, binding was notably more extensive at acidic pH (**Figure 3B**). Occupancy plots showed a higher number of SDS contacts with the scaffold at pH 3 compared to pH 5 over the course of simulations, indicating enhanced surfactant association under acidic conditions. Visual inspection further revealed that SDS clusters were stabilized around hydrophobic patches and aromatic residues lining the pores, suggesting a combination of hydrophobic and electrostatic interactions.

Complementary electrostatic surface mapping of the MS2 capsid dimers provided insight into differences in charge distributions across two pH conditions (**Figure S10**). At acidic pH, the outer shell of the capsid became increasingly positively charged compared to neutral or basic pH, driven by protonation of acidic residues. In contrast, the RNA-facing interior surface remained largely negative across all pH values. The protonation-induced shift towards positive surface charge at the outer capsid layer likely enhances electrostatic attraction to the negatively charged sulfate groups of SDS, promoting stronger and more penetrative binding at acidic pH.

Taken together, these results suggest that acidification not only increases capsid surface flexibility but also creates a favourable electrostatic landscape for SDS binding, concentrating surfactant molecules at structurally critical regions such as pores and dimer interfaces. This enhanced SDS accumulation may prime the capsid for subsequent destabilization and disassembly.

## 4. Discussion

### 4.1 Mechanistic Insights into SDS-Induced Disruption of the MS2 Capsid

The mechanism by which SDS disrupts viral capsids has long been hypothesized to involve surfactant-induced denaturation of viral proteins or global destabilization of viral architecture^35–37^. However, the specific structural cascade triggered by SDS—particularly related to viral inactivation mechanism at the molecular and sub-molecular level under different pH conditions—has remained poorly understood. This study integrates coarse-grained and all-atom MD simulations, together with experimental validation, to comprehensively resolve – for the first time - the SDS-driven destabilization mechanism of the MS2 bacteriophage. Experimentally, TEM and DLS analyses revealed that SDS exerts its virucidal effects in a highly pH-dependent manner. While MS2 particles largely retained their structural integrity at neutral pH, exposure to SDS at pH 2.5 triggered widespread capsid disassembly, as evidenced by the near-complete disappearance of intact particles and the appearance of sub-10□nm micellar-sized species. These findings suggest a pH-gated penetration mechanism, where acidic conditions facilitate SDS access to otherwise protected capsid interfaces. Notably, even modest pH increases (e.g., from 2.5 to 3.5) markedly reduced SDS efficacy, supporting the concept of a sharp transition threshold. This provides a mechanistic basis for the critical role of low pH in priming viral capsids for SDS-mediated disruption. Similar pH-dependent vulnerabilities have been reported for murine norovirus and MS2 bacteriophage in previous studies, where low pH exposure enhanced viral susceptibility to chemical treatments and destabilization, highlighting the broader importance of acidic environments in promoting antiviral efficacy^38,39^.

Based on simulations, the marked reduction in salt bridge interactions under acidic conditions indicates a weakening of the capsid’s electrostatic stabilization network, primarily driven by protonation of Glu and Asp residues. This loss of electrostatic integrity exposes basic residues (Lys, Arg), creating favourable binding sites for the negatively charged SDS headgroups. As a result, SDS micelles preferentially associate with structurally vulnerable regions, particularly inter-dimer clefts and pentameric and hexameric pores, with greater persistence and clustering observed at pH 2.5 compared to neutral pH. Occupancy maps, interaction lifetimes, and residue-level contact statistics all pointed to enhanced SDS binding with acidic residues at pH 2.5, particularly concentrated at hexameric pore openings—key mechanical weak points of the capsid. These findings align with prior studies emphasizing the essential role of salt bridges in maintaining capsid stability across diverse viral systems^40,41^ and mirror broader observations that low pH-mediated electrostatic repulsion can destabilize viral architectures and promote conformational changes^42^. Together, our results suggest that pH-driven salt bridge disruption acts as a critical molecular trigger, rendering the MS2 capsid more susceptible to SDS-mediated penetration and destabilization.

The MS2 capsid is known to naturally contain pores approximately 2-3 nm in diameter at its symmetry axes, allowing for molecular exchange between the capsid interior and the external environment. These pores, while typically rigid under neutral conditions, have been shown to exhibit pH- and ion-sensitive flexibility ^20,43^. In our CG simulations, SDS micelle binding led to significant expansion of hexameric pores, with occupied pores widening by an average of ∼3.7□Å under acidic conditions (pH 2.5), compared to minimal changes or slight contractions in unoccupied or neutral-pH systems. This expansion was corroborated by long-timescale simulations of a single hexameric pore, where SDS-bound pores expanded further to 45.4 Å versus 41.2 Å in the SDS-free system. These results suggest that SDS micelles exert mechanical pressure on the pore walls, actively driving dilation rather than merely binding passively. The enhanced flexibility and expansion of the pores under acidic conditions likely facilitate deeper SDS penetration and contributes to the destabilization of the capsid structure.

To elucidate how SDS micelle binding influences local capsid architecture, a multi-metric structural analysis was performed focusing on dimer–dimer interactions within the selected hexameric pore. COM distance measurements revealed that SDS binding increased separation between dimer pairs, where in some cases the SDS micelle density reached Δ ≈ 1.5 Å. Complementary tilt angle analysis showed that SDS binding promoted rotational “wobbling” of dimers, with an increase of up to 3.7° under SDS-free conditions and 7.9° in the SDS-bound system, indicating impaired rotational anchoring. Additionally, Rg analysis revealed an increase in internal flexibility, particularly adjacent to bound SDS micelles, suggesting local loosening of the dimer structure.

Collectively, these findings converge on a coherent mechanistic model of SDS-induced destabilization. Under acidic conditions, protonation of acidic residues disrupts stabilizing salt bridges, increasing capsid surface flexibility and exposing internal residues to surfactant penetration. SDS micelles preferentially bind to structurally vulnerable regions such as hexameric pores and inter-dimer clefts, initiating local perturbations characterized by pore dilation, increased inter-dimer spacing, rotational instability, and conformational loosening. These localized disruptions would likely propagate over longer timescales, weakening the global capsid framework and facilitating progressive disassembly. The consistency of these observations across experimental results, CG simulations, and all-atom models strongly supports the conclusion that SDS exploits latent structural vulnerabilities in the MS2 capsid, and that its enhanced virucidal efficacy at low pH arises from the synergistic effects of capsid softening and targeted micelle binding.

### 4.2 Toward safer and more effective antiviral surfactant strategies

Effective inactivation of non-enveloped viruses has traditionally relied on harsh, reactive chemicals, which are challenging to formulate for products intended for prolonged contact with human skin. Although SDS exhibits potent antiviral activity, its primary applications are outside of direct skin use due to its well-documented capacity to disrupt the skin barrier. Specifically, SDS can penetrate the *stratum corneum* through aqueous pores, increasing skin permeability and inducing irritation, particularly at concentrations above its critical micelle concentration. Mitigation strategies, such as the addition of humectants like glycerol, have been shown to reduce SDS-induced barrier perturbation, but the inherent irritancy of SDS still limits its suitability for extended skin contact in consumer formulations^44,45^. Understanding the molecular mechanisms underlying SDS-mediated viral inactivation, particularly its interactions with viral capsids and cellular membranes, is therefore critical for the rational design of next-generation antiviral agents. By dissecting the physicochemical features that enable SDS to destabilize robust non-enveloped viruses while also recognizing its limitations in terms of biocompatibility, it becomes possible to inform the development of alternative surfactant systems, formulations, or targeted delivery strategies that maintain virucidal efficacy while minimizing skin irritation.

Our findings strongly align with previous studies demonstrating the virucidal efficacy of SDS against a broad range of viruses, while offering new mechanistic insights into its mode of action. Earlier work primarily attributed SDS-mediated inactivation to general protein denaturation and membrane solubilization mechanisms^14,46,47^. However, direct structural evidence at the viral capsid level has been limited. Here, by combining CG and all-atom simulations with experimental validation, SDS was shown to preferentially target hexameric pores and inter-dimer clefts—sites of intrinsic mechanical vulnerability in the MS2 capsid— where binding initiates local pore dilation, inter-dimer separation, and conformational loosening. These localized structural perturbations propagate over time, weakening the capsid framework and promoting progressive disassembly. In doing so, our study provides a detailed molecular pathway for SDS-driven destabilization, extending beyond previous models of nonspecific surfactant-induced disruption. Notably, the mechanistic principles uncovered here may also apply more broadly to other non-enveloped viruses that share similar icosahedral architectures and pore-like surface features. These insights lay the groundwork for the rational design of novel surfactant-based antiviral strategies that optimize virucidal efficacy while minimizing collateral damage to human tissues. A rational approach to improving surfactant biocompatibility could involve modifying molecular features that govern skin barrier disruption. One strategy is to adjust the hydrophobic tail length or to introduce hydrophilic spacers between the sulfate headgroup and alkyl chain, thereby reducing deep penetration into the stratum corneum. Alternatively, replacing the sulfate headgroup with milder functional groups—such as carboxylates or zwitterionic moieties— may mitigate surfactant-induced barrier perturbation. Notably, surfactants like sodium laureth sulfate (SLES) and certain betaine derivatives have demonstrated reduced irritancy while maintaining strong surface activity^2,48^.

Beyond direct modification of SDS, alternative surfactant classes offer promising avenues for antiviral formulation development. Zwitterionic surfactants, sugar-based surfactants (e.g., alkyl polyglucosides), and synthetic peptide-mimetic surfactants combine desirable surface activity with enhanced biocompatibility and represent attractive candidates for future investigation. These alternatives could maintain virucidal efficacy by exploiting tailored electrostatic and hydrophobic interactions with viral capsids, while minimizing adverse effects on mammalian skin barriers. Furthermore, recent studies have explored the combination of surfactants with organic acids, such as citric and succinic acids, to enhance virucidal activity while potentially mitigating skin irritation^14,49,50^. Although the precise mechanisms by which organic acids contribute to virus destabilization remain speculative, proposed mechanisms include pH-induced structural weakening and direct interactions with the genome. Future investigations dissecting the synergistic actions between safer surfactants and organic acids may enable the rational design of next-generation antiviral formulations that optimize both efficacy and skin compatibility.

Overall, elucidating the molecular principles underlying SDS-mediated capsid destabilization provides a critical foundation for the design of safer and more effective antiviral surfactants. Future work combining molecular dynamics simulations with experimental screening of modified surfactant libraries may accelerate the discovery of optimized agents suitable for topical or consumer use.

## 5. Conclusions

In this study, a combined experimental approaches with CG and all-atom molecular dynamics simulations to dissect the mechanism of SDS-mediated disruption of the MS2 bacteriophage capsid was explored. Experimental analyses using TEM and DLS confirmed that SDS efficacy is highly pH-dependent, with significant capsid disassembly occurring at acidic pH but not at neutral conditions. To mechanistically resolve these observations, molecular simulations were employed across multiple scales: coarse-grained models captured long-timescale capsid responses to SDS binding, while all-atom scaffold models provided residue-level insights into SDS–protein interactions at pore and dimer interfaces. Together, these complementary methods enabled a detailed characterization of the structural and electrostatic factors underlying SDS-driven capsid destabilization.

Our findings reveal that SDS destabilizes the MS2 capsid through a multifaceted mechanism involving pH-triggered salt bridge disruption, micelle binding at structurally vulnerable regions such as hexameric pores and dimer interfaces, pore expansion, and localized dimer deformation. Under acidic conditions, protonation of acidic residues weakens the capsid’s electrostatic network, enhancing SDS accessibility and promoting deeper surfactant penetration. SDS micelle binding induces pore dilation, increases dimer separation, promotes rotational instability, and loosens internal dimer structure — changes that collectively undermine capsid integrity and likely facilitate progressive disassembly. These results provide a coherent molecular framework for understanding SDS-mediated virucidal action and highlight the importance of environmental factors such as pH in modulating viral susceptibility to surfactant disruption.

## Acknowledgements

The computational work for this study was carried out using resources from the National Supercomputing Centre, Singapore (https://www.nscc.sg), and the internal high-performance computing cluster of Procter & Gamble (P&G).

## Author Contributions

Conceptualization: PJB, CSV, SJB, JL, CMM

Methodology: SM, JKM, CC, HC, PJB, CSV, SJF, CMM

Software: SM, JKM

Validation: SM, JKM

Formal analysis: SM, JKM, MLYK, GL

Investigation: SM, JKM

Resources: PJB, SJF, CSV, CMM, CC

Data curation: SM, JKM

Writing—original draft: SM, JKM

Writing – Review and Editing: SM, JKM, PJB, SJF, CMM

Visualization: SM, JKM, PJB, SJF, CSV, CC, MLYK

Supervision: PJB, CSV, SJF, CC, CMM

Project administration: PJB, CSV, SJF, CC, CMM

## Data Availability

Correspondence and requests for materials should be addressed to Peter J. Bond.

