## Supplementary information for "A Multiscale Framework for Uncovering Surfactant Mediated Viral Capsid Disruption"

### **Supplementary Methods**

### **Maturation protein exclusion**

To mechanistically dissect the specific effects of SDS on capsid architecture, we performed simulations on MS2 particles excluding the MP. While the MP plays a key role in host recognition and introduces structural asymmetry in the capsid, our preliminary simulations with and without the MP revealed that SDS preferentially interacts with capsid pore regions and dimer interfaces, rather than with the MP itself (Figure below). Additionally, our previous comprehensive study demonstrated no substantial differences in pore expansion, overall capsid dynamics due to the presence or absence of the MP^22^. In addition, distribution of salt bridges at two pH conditions tested was following the same trend independently of the presence or absence of MP (Figure below). Moreover, supporting experimental data from DLS and Cryo-EM further indicated complete disassembly of MS2 capsids into sub-10 nm fragments, suggesting SDS-induced disruption primarily targets structural integrity via key capsid interfaces, rather than via the MP-mediated attachment pathway. Thus, to clearly define and generalize the biophysical mechanism underlying SDS-driven capsid disruption, we modelled the capsid in its symmetric, MP-free form. This approach allowed us to explicitly investigate pore dilation, salt bridge destabilization, and dimer interface separation as core mechanisms driving viral inactivation.

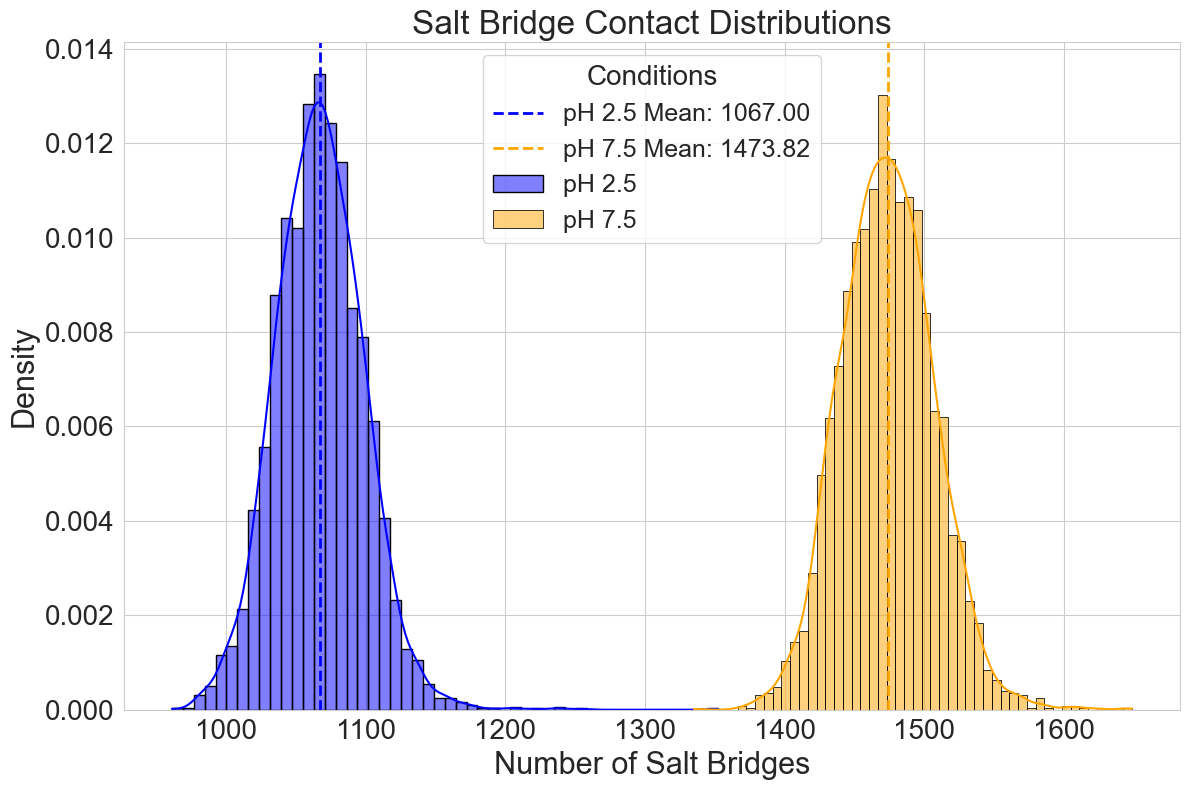

**Supplementary Figure. Salt-bridge contact analysis of MS2 capsid with maturation protein at different pH conditions.** Distribution of salt bridge contacts between acidic (Glu/Asp) and basic (Lys/Arg) residues over the last 2 μs of simulation. The number of salt bridges is significantly lower at pH 2.5 compared to pH 7.5, consistent with the expected protonation of Glu and Asp residues at low pH, reducing electrostatic interactions.

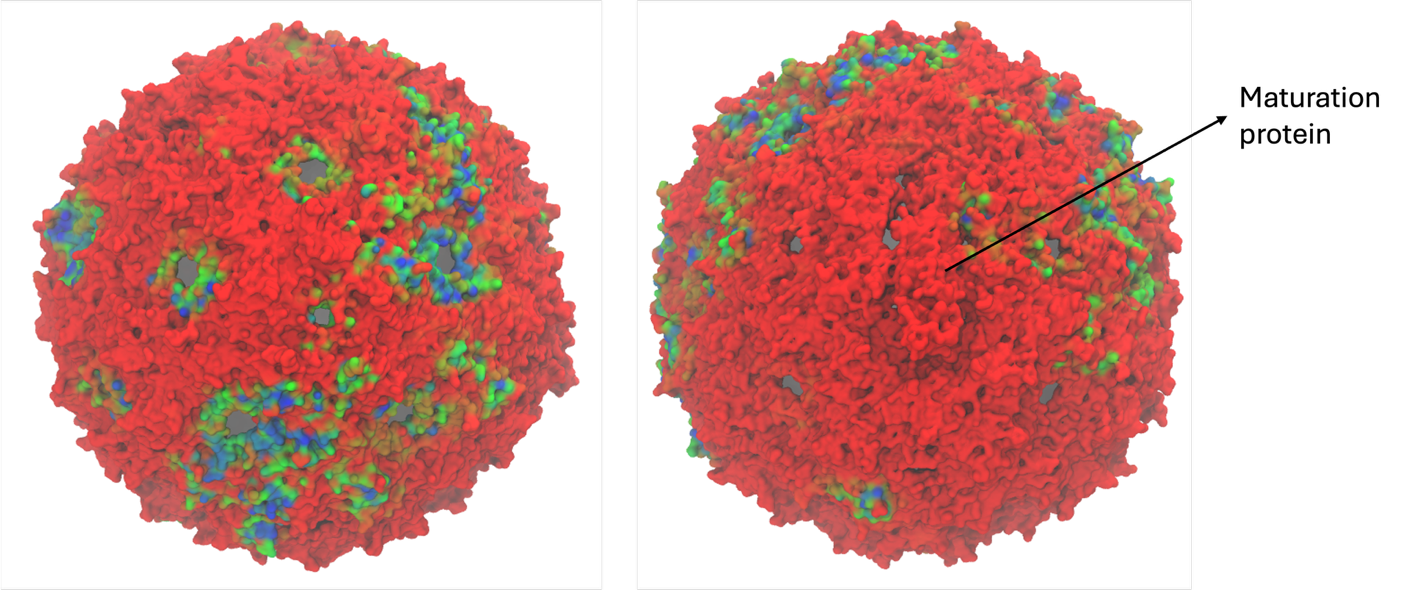

**Supplementary Figure.** Surface occupancy maps showing SDS accumulation on the MS2 capsid with maturation protein. Red regions denote low occupancy; blue and green regions denote higher occupancy.

### **Calibration of elastic network parameters to match All-Atom dynamics**

CG models, such as those based on the Martini force field, often require additional restraints to maintain the structural integrity of biomolecular assemblies due to their reduced degrees of freedom. Elastic networks, which apply harmonic restraints between selected atoms, are widely used to stabilize the tertiary and quaternary structures of proteins and protein complexes, compensating for missing atomistic detail and ensuring biologically realistic dynamics. To reliably capture the structural dynamics of the MS2 bacteriophage capsid at the coarse-grained level, the necessity of including an elastic network was evaluated and then the parameters were optimized by calibrating against reference all-atom simulations. Simulations performed without an elastic network showed pronounced structural instability, with RMSD values surpassing 1.2 nm and failing to stabilize over timescale of 500 ns. Introducing an elastic network restricted to secondary structure elements significantly enhanced capsid stability, yielding lower RMSD values more closely aligned with those from the all-atom reference simulations, and demonstrating a plateau indicative of structural equilibration (**Figure S1A**). However, minor deviations remained, suggesting room for further parameter refinement. To systematically optimize the elastic network, a range of force constants (Fc) were tested. Results indicated that increasing Fc from 500 to 3,000 kJ mol^−1^ nm^−2^ progressively improved structural fidelity, markedly reducing RMSD fluctuations (**Figure S1B**). Beyond an Fc of 3,000 kJ mol^−1^ nm^−2^, additional increases in force constant yielded negligible RMSD improvement, indicating saturation. Thus, an Fc value of 3,000 kJ mol^−1^ nm^−2^ was determined to offer the most accurate representation for CG model, reproducing dynamics observed in all-atom simulations.

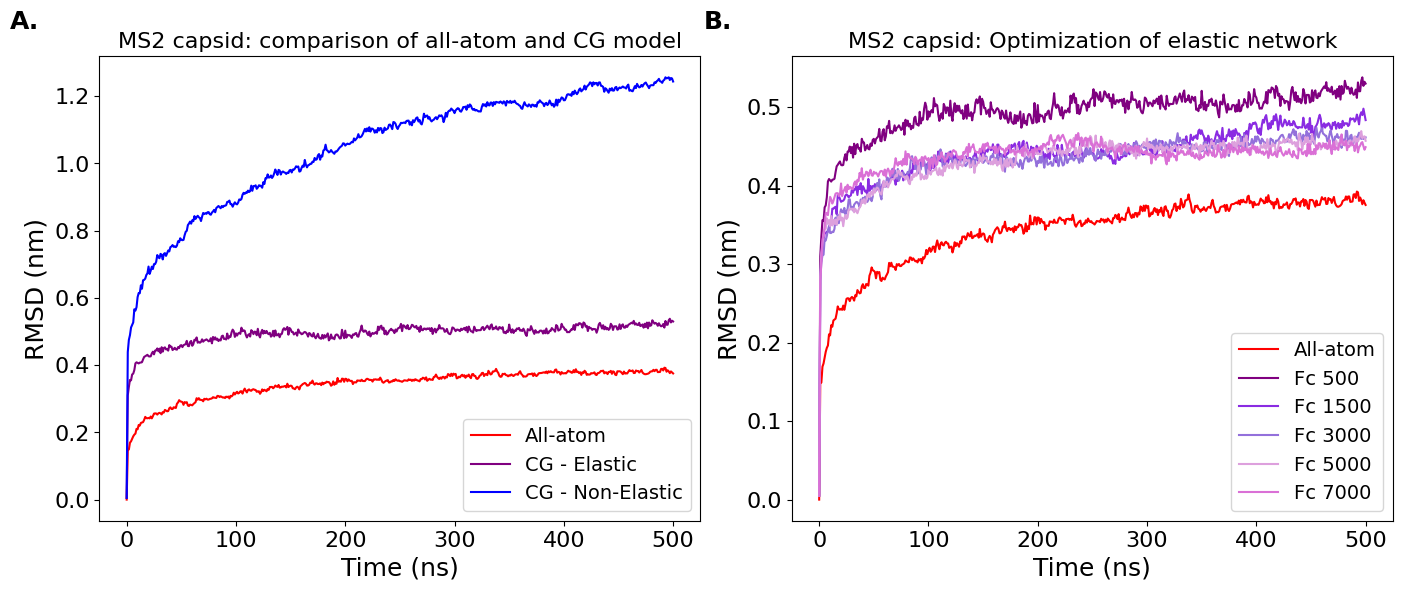

**Figure S1. Backbone RMSD analysis of MS2 capsid models. A.** Comparison of all-atom and CG models of the MS2 capsid. The all-atom model (red) shows higher stability compared to the CG models, with the elastic network (purple) being more stable than the non-elastic network (blue). **B.** Optimization of the elastic network in the CG model of the MS2 capsid. Different force constants (Fc) are tested, with higher force constants (e.g., Fc 7,000 kJ mol^−1^ nm^−2^, orchid) showing improved stability compared to lower force constants (e.g., Fc 500 kJ mol^−1^ nm^−2^, purple). RMSD values are plotted as a function of simulation time (in nanoseconds).

**Limitations**

While this study provides detailed insights into SDS-mediated destabilization of the MS2 capsid, several limitations must be acknowledged. First, the use of the Martini coarse-grained force field, while computationally efficient, may introduce artifacts such as over stabilized non-bonded interactions, which can lead to exaggerated SDS aggregation and potentially underrepresent the intrinsic stability of the protein interfaces. Second, our CG models do not include the viral RNA genome, which is known to contribute significantly to capsid stability. The absence of this internal structural component may partially explain why complete capsid collapse was not observed, even under high SDS concentrations and extended simulation timescales. Finally, despite testing a variety of perturbative conditions—including 50% SDS concentration, Martini 3 parameter scaling, and simulations extended up to 44 microseconds—the capsid remained partially intact, highlighting the combined impact of missing internal forces and force field limitations. Future work could benefit from the inclusion of RNA, improved parameter sets such as those in Martini 3.2, and the integration of longer all-atom simulations to validate and refine CG MD predictions. Additionally, exploring SDS activity in combination with other destabilizing agents may provide a more complete picture of synergistic viral inactivation mechanisms.

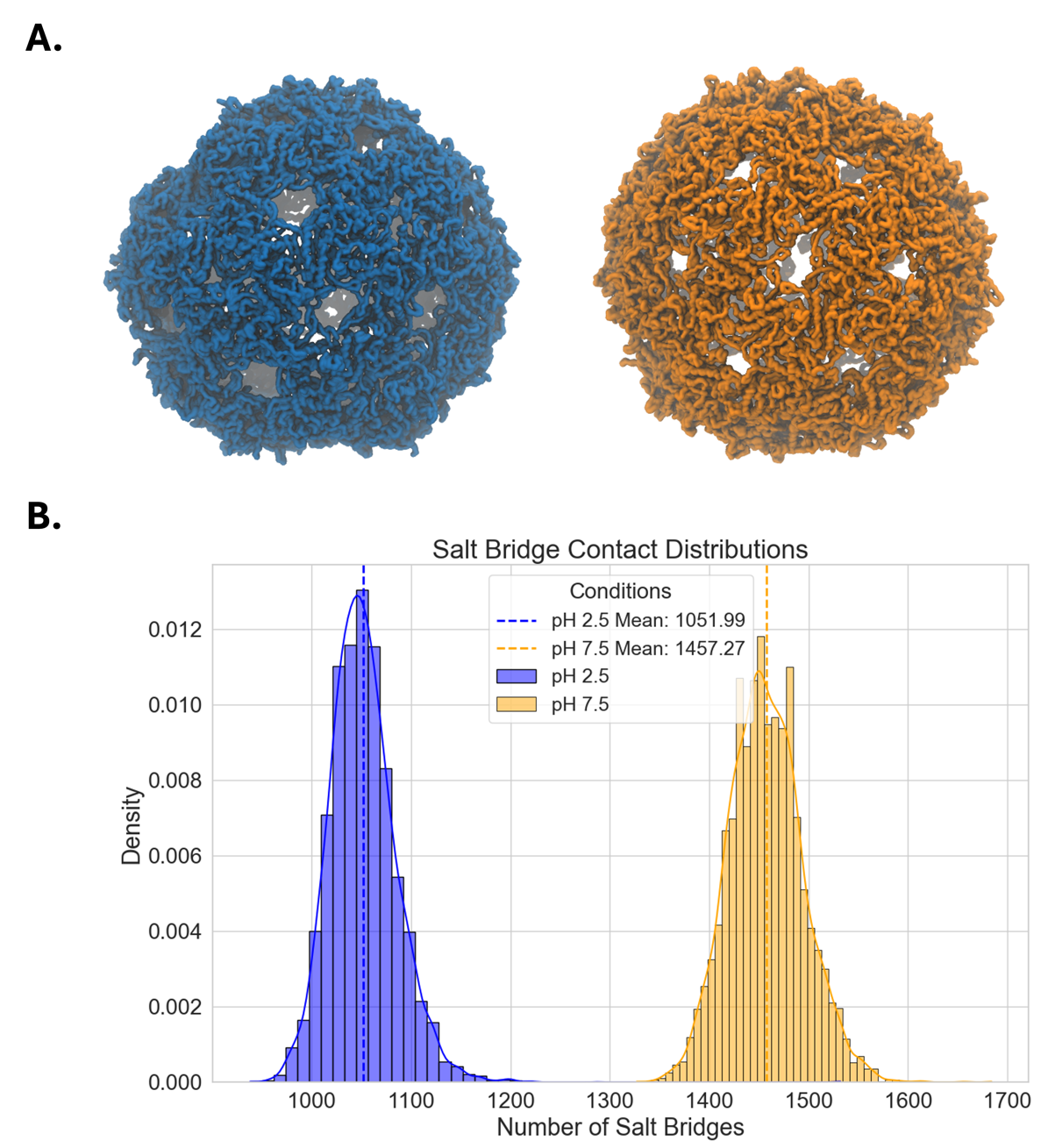

**Figure S2. Structural and electrostatic analysis of MS2 capsid at different pH conditions.
A.** CG representation of the MS2 capsid at pH 2.5, showing structural organization under acidic conditions (left); CG representation of the MS2 capsid at pH 7.5, illustrating capsid shape under neutral conditions (right). **B.** Distribution of salt bridge contacts between acidic (Glu/Asp) and basic (Lys/Arg) residues over the last 2 μs of simulation. The number of salt bridges is significantly lower at pH 2.5 compared to pH 7.5, consistent with the expected protonation of Glu and Asp residues at low pH, reducing electrostatic interactions.

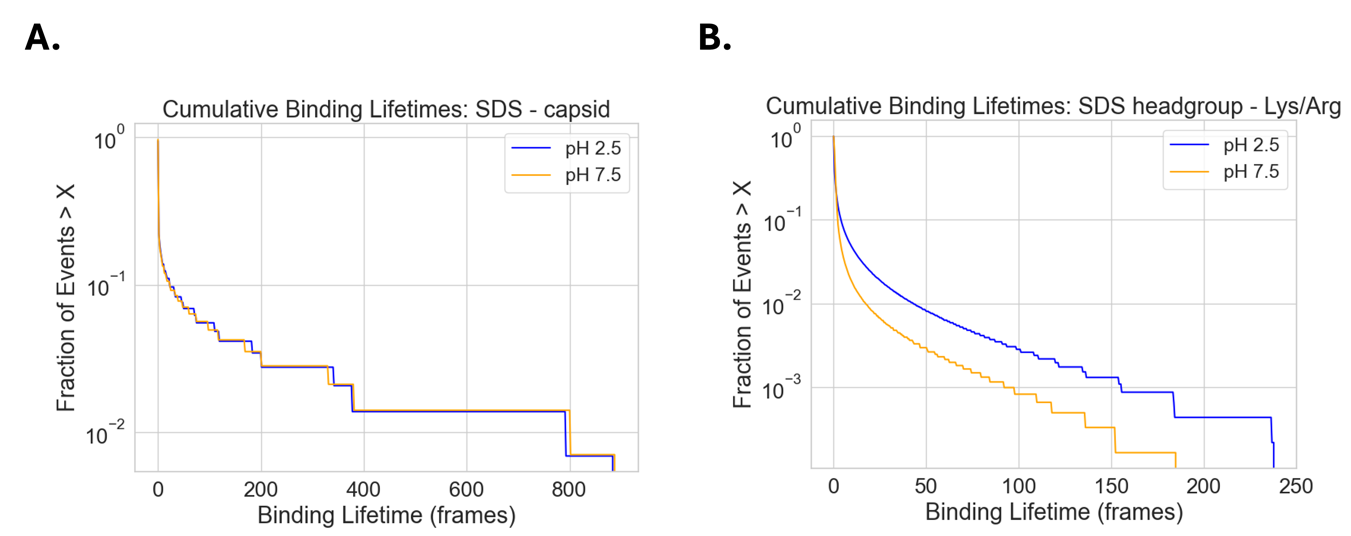

**Figure S3. Binding lifetime analysis between SDS and MS2 capsid at different pH conditions.** **A.** Binding lifetimes between SDS molecules and the entire MS2 capsid (within 6 Å) are similar at both pH 2.5 and pH 7.5. **B.** Binding lifetimes between the SDS headgroup (SO₃⁻) and positively charged Lys/Arg residues (within 6 Å) show distinct behaviour: although at pH 2.5 these interactions occur more frequently (see Figure S6), and they are significantly longer-lived than those at pH 7.5.

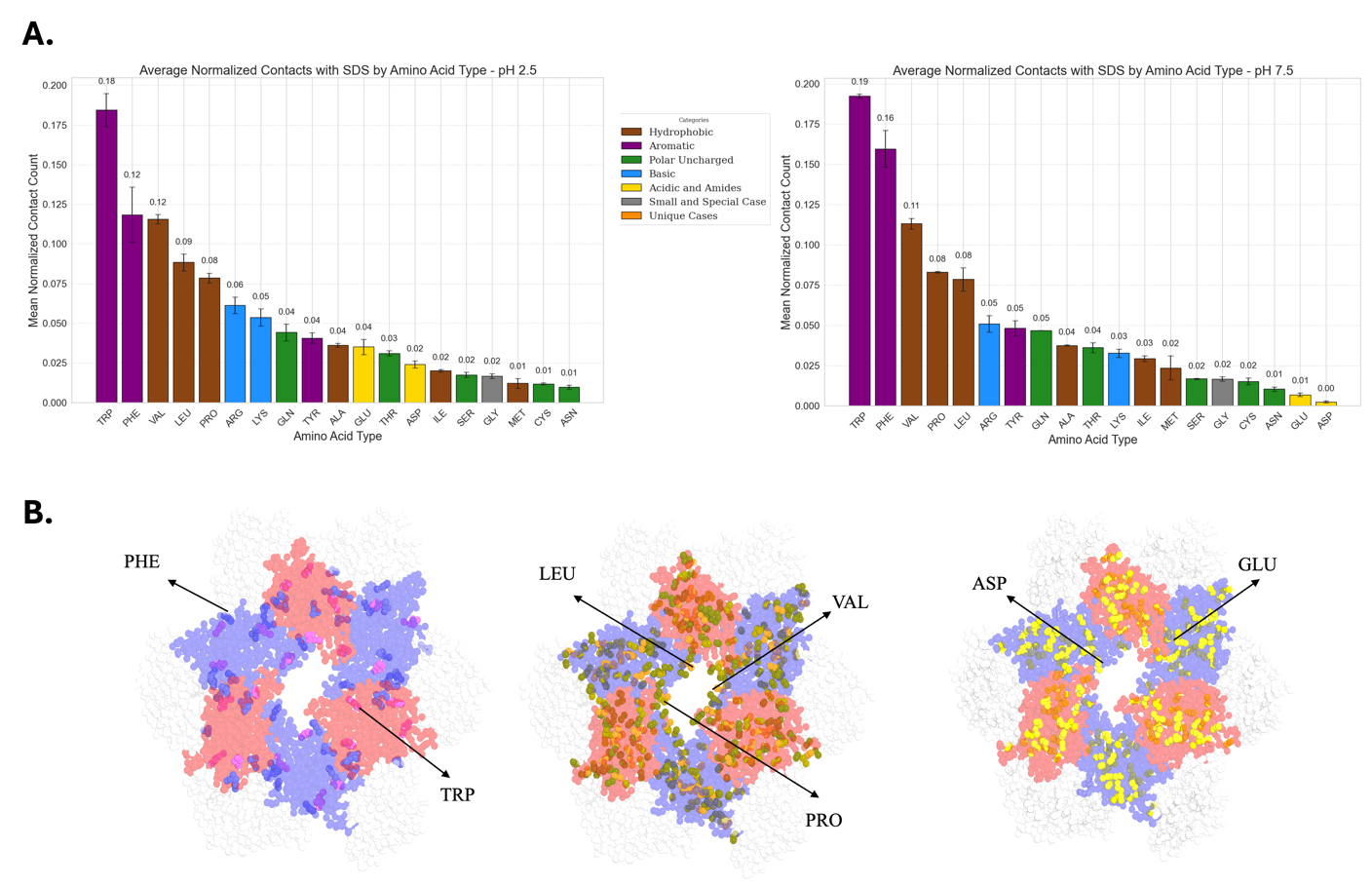
**Figure S4. Comparison of SDS–protein interactions at acidic and neutral pH conditions.**
**A.** Bar plots showing the average normalized contact frequency between SDS molecules and amino acid types in the MS2 capsid, calculated over the last 2 μs at **pH 2.5** (left) and **pH 7.5** (right). Aromatic residues (Trp, Phe) and hydrophobic residues (Val, Leu, Pro) exhibit the strongest interactions with SDS. Notably, Glu, Asp show higher contact frequencies at **pH 2.5**, suggesting enhanced SDS accessibility to acidic surface patches due to protonation differences between the two pH conditions. **B.** PyMOL-rendered illustrations of the MS2 capsid highlighting locations of Trp and Phe residues (shades of purple); Pro, Leu, and Val residues (shades of brown); and Glu/Asp residues (shades of yellow) in three representative hexamer units. Aromatic residues cluster prominently at the dimer interface regions, while acidic and hydrophobic residues cluster at the pore regions, likely serving as key hotspots for SDS association.

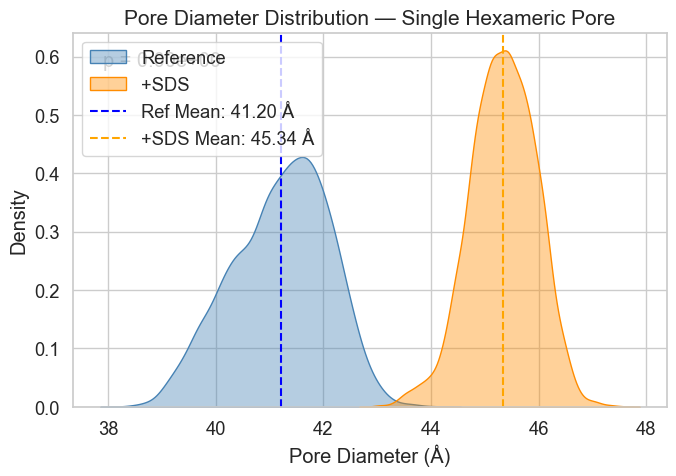

**Figure S5. Pore diameter comparison in the absence and presence of SDS. – CG simulations.** Comparison of pore diameter distributions for a representative hexameric pore under reference (blue) and SDS-bound (orange) conditions. Kernel density estimates show a shift toward larger pore diameters upon SDS binding. The mean pore diameter increased from 41.2 Å (reference-SDS_free_) to 45.3 Å (+SDS). A statistical comparison using a two-sample test yields a p-value of 0.00, indicating a significant difference between the two distributions.

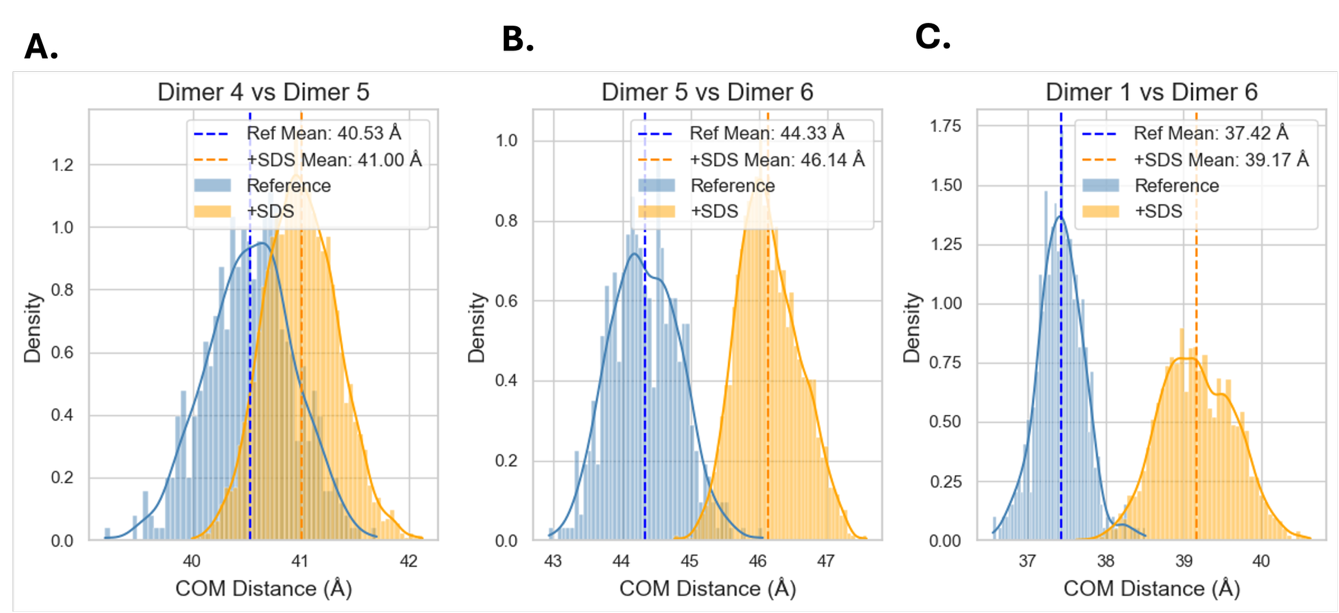

**Figure S6**. **COM distribution comparison in the absence and presence of SDS – CG simulations.** COM distance distributions for selected dimer pairs in the hexameric pore under reference (blue) and SDS-bound (orange) conditions. Each panel shows a comparison for one dimer pair: **A.** Dimer 4 vs Dimer 5, **B.** Dimer 5 vs Dimer 6, and **C.** Dimer 1 vs Dimer 6. Kernel density estimates and histograms illustrate shifts toward larger COM distances upon SDS binding. Mean COM distances for each condition are indicated by dashed vertical lines. A consistent increase in COM separation upon SDS interaction suggests pore expansion or dimer destabilization.

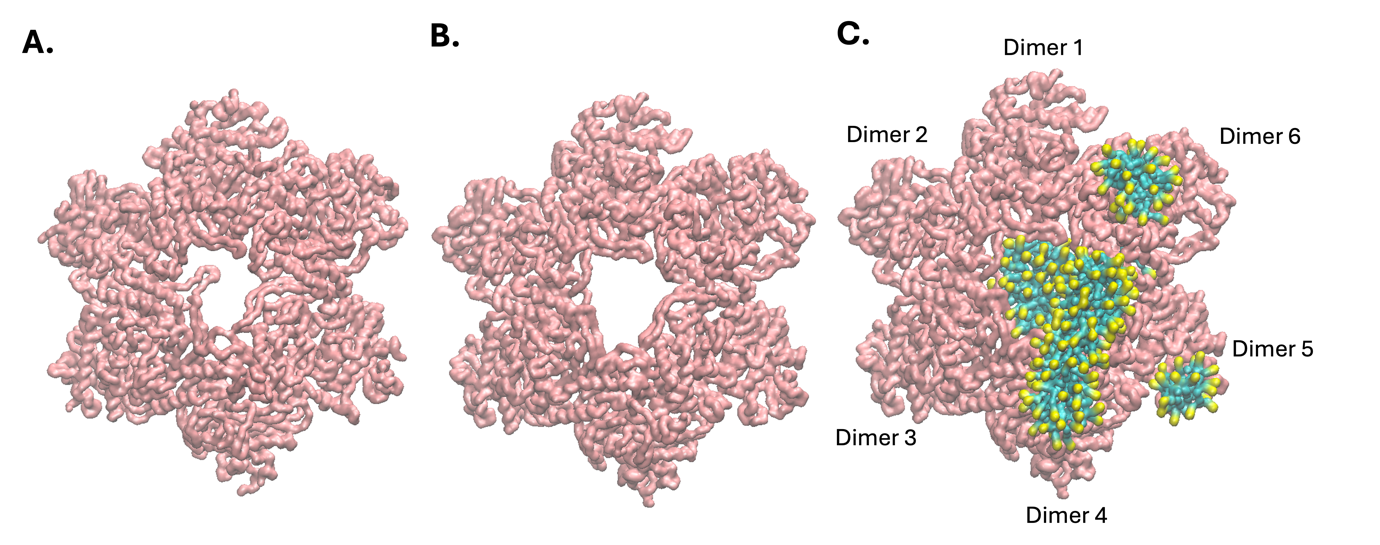

**Figure S7. Long-timescale simulation of SDS-bound hexamer reveals progressive pore expansion and dimer perturbation. A.** Structure of the MS2 hexameric pore after 10 μs of CG simulation at pH 2.5 in the absence of SDS, showing preserved architecture. **B.** Same hexameric pore after a 5 μs simulation with SDS micelle present (SDS removed for clarity), displaying notable pore expansion and interface loosening. **C.** Final structure from the SDS-bound simulation with SDS molecules shown (yellow headgroups, blue tails), highlighting clustering at inter-dimer clefts. Dimer units (Dimer 1–6) are labelled for reference to downstream structural analyses.

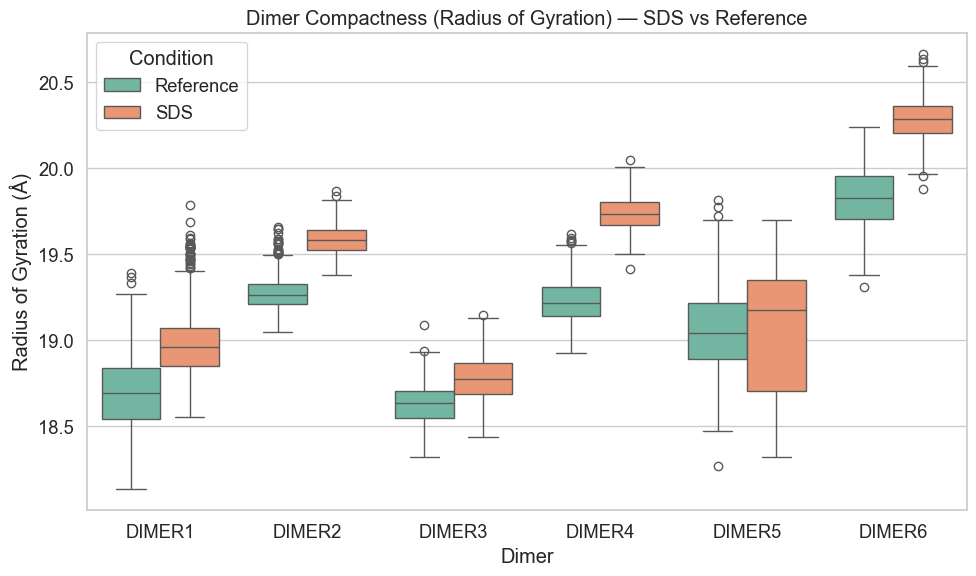

**Figure S8.** **Dimer Compactness – CG simulations.** Boxplots showing the radius of gyration for each capsid dimer (Dimer 1–Dimer 6) under reference conditions (green) and after SDS treatment (orange). A general increase in Rg is observed upon SDS binding, indicating a loosening or destabilization effect across several dimers. Each box represents the interquartile range, with the line indicating the median.

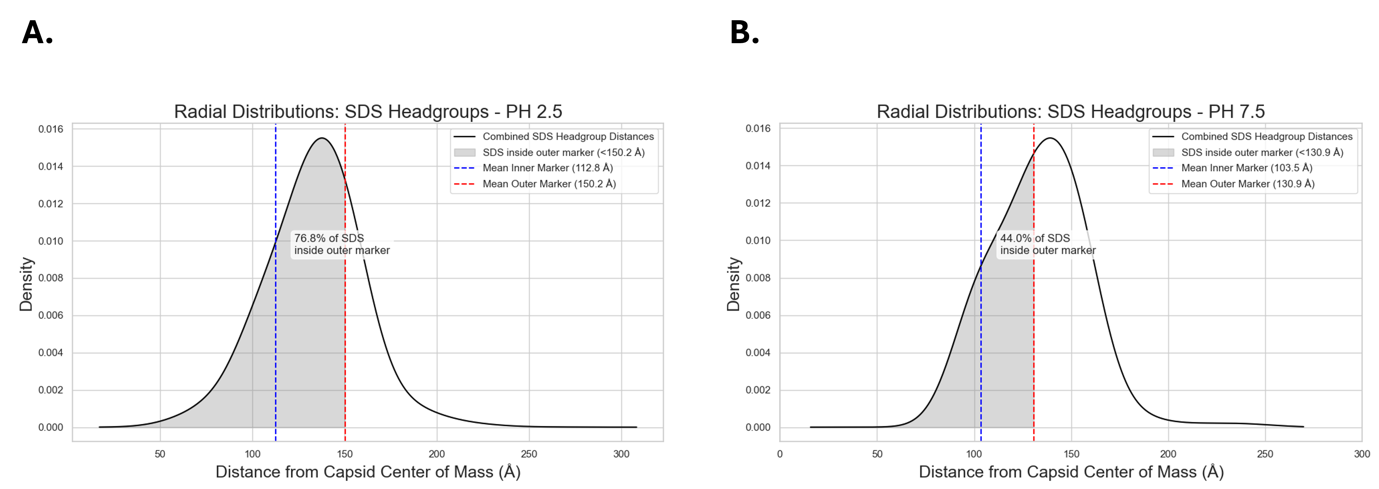

**Figure S9. Radial distribution of SDS headgroup distances from the capsid COM at pH 2.5 (A) and pH 7.5 (B) – CG simulations.** The blue dashed line represents the average distance to the inner capsid marker (closest residue to the center), and the red dashed line indicates the outer capsid marker (furthest residue). Shaded areas highlight SDS headgroups located within the outer capsid boundary. At pH 2.5, **76.8%** of SDS molecules are localized inside the outer boundary, compared to **44.0%** at pH 7.5, indicating deeper SDS penetration into the capsid at acidic pH. This suggests increased capsid flexibility and expansion at low pH, facilitating enhanced SDS burial within the viral structure.

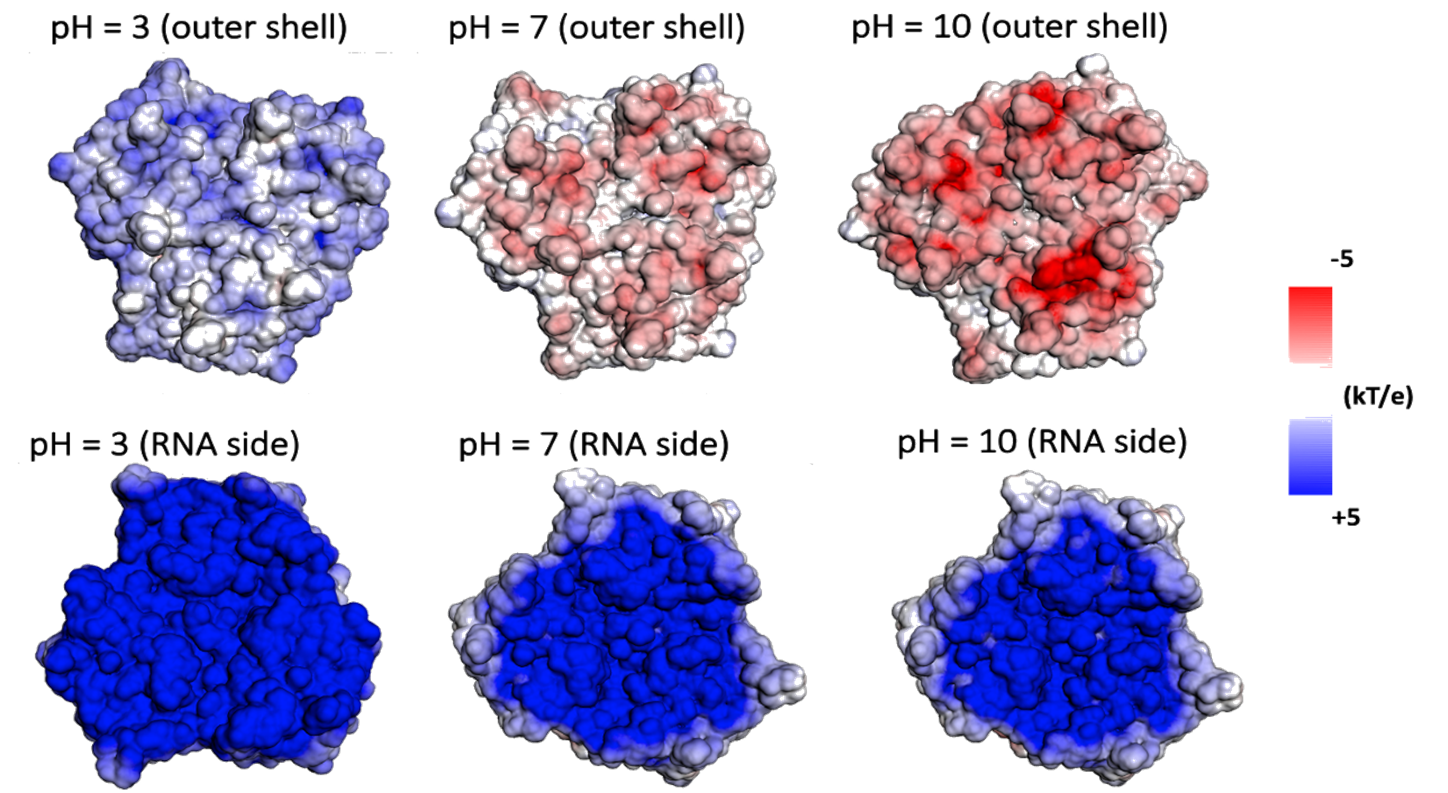

**Figure S10. Electrostatic potential maps of MS2 capsid trimer under varying pH conditions – All-atom simulations.** Surface representations of the MS2 capsid calculated using APBS electrostatics, shown from the outer (top row) and RNA-facing (bottom row) sides at pH 2, 7, and 10. The capsid becomes increasingly positively charged at acidic pH, particularly on the outer surface, facilitating electrostatic attraction with the negatively charged headgroups of SDS molecules. The inner surface remains predominantly neutral across all conditions, suggesting SDS primarily interacts with the exterior regions of the capsid.
